# Isolating Compounds that Inhibit EV 71 Virus

**DOI:** 10.1101/2020.07.30.229815

**Authors:** Giridhar Murali, Rishabh Kejriwal, David Olson, Lauren Alexandrescu, Simon White

**Affiliations:** University of Connecticut

## Abstract

Enterovirus 71, or EV 71, is responsible for causing Hand, Foot, and Mouth disease in humans. In particular, it is especially deadly when children and small infants are exposed. The objective of this research paper is to address the possibility of a novel antiviral drug that can be used once infection of EV 71 has occurred. The methods for this research include transformation of *E. coli* with the genetic information from Enterovirus 71, growth of the *E. coli* colonies in the lab setting, 2C protein purification, and ATPase assays with drug testing. Of the 364 drugs tested in the ATPase assay, a combination of two of them (Mitrofudil and N6-Benzyladenosine) indicated a stoppage in activity of ATPase, signaling no further activity of the enzyme and viral proliferation.

## Introduction

Human Enterovirus 71 (EV71) is a small, non-enveloped, icosahedral virus that belongs to the Enterovirus genus within the family Picornaviridae (Guan *et al*, 2019). It is a virus that infects humans and is the causative agent of hand, foot, and mouth disease, which causes fever and flu-like symptoms, mouth sores, and skin rash (Belazarian L, Lorenzo ME, 2017). However, severe cases can lead to paralysis or encephalitis, which can prove to be fatal in the patient. There have been breakouts of hand, foot, and mouth disease throughout the world, with the most prominent cases occurring in Eastern Asia in populations of infants and small children (Wang *et al,* 2019). During the viral lifecycle it relies on a viral protein known as 2C, which is responsible for viral replication and proliferation. 2C has ATPase activity that it needs for function (Yun, 2010).

It is vital to study EV 71 reproduction in order to facilitate the development of an effective antiviral agent in the future. Currently, there exists no specific treatment protocol for hand, foot, and mouth disease (Liu, 2010). Many physicians prescribe a topical oral anesthetic to aid in relieving the pain associated with mouth sores, but it cannot accelerate the clearance of the disease from the seven to ten-day infection period (Berman, 2017). A vaccine against EV 71 is presently available in China, but others are still being actively researched and studied (Dong *et al*, 2019).

Based on these factors, it is important to understand the replication process of EV 71 through the 2C protein, which is one of the most highly conserved proteins containing both ATPase and RNA helicase activity, in order to find an antiviral compound (Cardosa, 2007). The objective of this research thesis was to identify a novel antiviral compound that can be used once infection of EV 71 has occurred. The methods for this research include transformation of *E. coli* with the 2C gene from Enterovirus 71, growth of the *E. coli* colonies in the lab setting, 2C protein purification, and ATPase assays with drug testing. This project was able to find two antiviral compounds that were able to limit the activity of the ATPase activity of the 2C protein, which can be further developed into commercial antivirals in the future for patients suffering from hand, foot, and mouth disease.

## Materials and Methods

### Transformation of *E. coli* with the genetic information from Enterovirus 71

To begin, *E. coli* plates were grown which had been transformed with the genetic information, specifically the viral replication proteins, from EV 71. This was done by microwaving a plate agar bottle. The lid was loosened and the microwave was set for a time limit of 5 minutes. While heating, it was important to take out the bottle every 15 seconds to stir. After heating, the lid of the bottle was tightened and let to cool. The Bunsen Burner was turned on and a 50ml Falcon tube was taken out, and the following was poured: 50ml of agar (LB agar, 32g per L of H2O), 50 microliters of Ampicillin at a 25 mg/ml concentration, and 50 microliters of Chloramphenicol at a 25 mg/ml concentration. The reasoning behind choosing Ampicillin and Chloramphenicol was to select the colonies that had genetically altered plasmids which were resistant to these antibiotics in order to grow up enough protein for the purification process (Tu, 2007). This was useful since these plasmids could be grown with conferred antibiotic resistance, which ensured that the 2C protein could be extracted. Two plates were made, so 25 ml of solution was poured into one plate and 25 ml of solution into another. If *E. coli* samples were not already readily available, transformation needed to be done. *E. coli,* specifically the Arctic Express (DE3) Competent Cell strain, was used in this process for producing enough protein for purification. This strain was used over others since it had the capacity for growth in a relatively short time, could be grown at lower temperatures, and handle higher toxicity levels (Herrero *et al,* 2003). For transformation, a bucket of ice was taken and the EV 71 2C was acquired, which was amplified through Polymerase Chain Reaction (PCR), from the fridge. Also, purple PCR tubes were acquired which contained the Arctic Express (DE3) Competent *E. coli* cell pellets. The PCR tubes were then labeled. The hot water bath was turned on to 44 degrees C and the power button was hit. Ten microliters of each EV 71 sample was added into the respective PCR tubes. The tubes were flicked and placed back in ice for 30 min. After 30 minutes, the two tubes were heat shocked by being placed into the heated water bath for 30 seconds. The Petri dishes were labeled. 450 microliters of SOC media (composed of 2% Tryptone, 0.5% Yeast Extract, 10mM NaCl, 2.5mM KCl, 10mM MgCl2, and 20mM Glucose) was added to the small PCR tubes at a maximum volume of 0.2ml. The purpose of the SOC media was to provide nutrients for the *E. coli* in order to grow colonies. 100 microliters of sample from the small PCR tube was added to the agar plate alongside 20 microliters for each of the four overnight tubes, which were 50ml Falcon tubes that contained SOC media, antibiotics, and a picked colony of *E. coli* to grow overnight. The plate stick was put in the flame to sterilize. This was repeated once more. The stick was inverted and streaked across the plate. Overnights and plates were placed into a shaker to incubate overnight. All tubes and plates were ensured to be labeled.

### Growth of *E. coli* colonies in the lab setting

The four overnights were taken out with the bacteria/virus alongside the SOC media at a 20 ml volume. Four large 2.8 L growing flasks were filled with 2XYT LB media (composed of 24g of Tryptone, 15g of Yeast extract, 3g of Glucose, and 7.5g of NaCl) at a volume of 1.5L that has been pre prepared. 1 ml of Ampicillin was added at a 25mg/ml concentration as well as 1 ml of Chloramphenicol at a 25 mg/ml concentration to the big flask. The overnight was taken and poured into the growing flask. Aluminum foil was placed on the lid and the flask was put back into the shaker. It was incubated for 2 hrs. at a temperature of 37 degrees Celsius. A nanodrop was conducted by taking out 1 ml of sample and 1 ml of original growth media. The nanodrop machine was used to set the original growth media as a blank to compare with the new sample reading. A 0.5 OD (Optical Density) value from the reader was the OD value that was important to note in the samples. This OD value signified that *E. coli* growth was sufficient in order to have enough cells to purify the 2C protein (McMinn, 2001). 1 ml of IPTG was added to the growing flask. IPTG was used to induce protein expression under control of the lac operon in *E. coli* (Chan *et al,* 2003). The flask was placed back into the shaker and the incubation temperature was changed to 30 degrees Celsius. The flasks were incubated for an additional 6 hours. The liquid was poured into 2 small plastic centrifuge containers. The masses were balanced out. They were placed into the centrifuge for 15 minutes. The supernatant was poured out and the cell pellet was stored at −80 degrees Celsius.

### 2C protein purification

A 50 ml of Buffer A (composed of 0.02M of Bis-Tris and 0.2M of NaCl at pH=7.0 at 1L volume) was placed into a 50 ml blue-top Falcon tube.1 protease inhibitor tablet was added, and dissolved. Buffer A was placed back in the fridge. The tube was labeled as: Buffer A + protease inhibitor. 10 ml of the solution was placed into each pellet. It was mixed thoroughly. Two Oakridge centrifuge tubes were taken out and placed on ice. The lid was removed off of one of them. The Constant One-shot system was utilized in order to lyse the cells. One-shot was important to break open the cells for extracting and purifying the 2C protein. In order to operate the Constant One-shot system, a silver cap with a lid was needed. A total volume of 7ml was lysed for each blast. The cell lysate was placed parallel with the opening and pushed down. The locking mechanism was aligned by matching the dots. A metallic rod was placed to secure the sample and other components. It was removed once completed and pushed off the liquid. A finger was placed on the anti-foam device and poured into the tube. It was kept on ice. After this, a small beaker was acquired and balanced out with the other tube. The tube was balanced with distilled water. The speed on the centrifuge was changed to 25,000 xG and spun for 30 minutes. This step was essential for creating a cell pellet with supernatant on the top (Goh *et al,* 2006). The cell debris was to be found in the pellet, and the supernatant was further purified. This was proceeded via the AKTA system control with 0.22M Buffer A and Buffer B (composed of 0.01M Bis-Tris, 0.1M NaCl, and 0.25M Imidazole at pH=7.0 at 0.5L volume) at a flow rate of 5 ml/min with a Nickel column. A Nickel column was used for ensuring the 2C protein can adhere to the column and then would be eluted off at a specific concentration, which can be determined via SDS-PAGE (Chatproedprai *et al*, 2010).

Manual 1 was pressed on the system to change the flow rate to 5 ml/min. The blue button was pressed to make the sample value green. It was switched off when the volume reached the 10 ml mark. It was turned back to Buffer B at that point for another 10ml. Next, it was switched to Buffer A and given 10 ml. Finally, it was switched to a weaker value to provide 20 ml. The rotor was filled up with empty tubes in the meantime. The stop button was pressed once 20 ml of weak value were added (square button). The tube was pushed over to (1), the starting position. To filter the samples, the tubes were taken out of the centrifuge. An ice box was bought with a 100ml bottle inside. The filter apparatus was set up by placing filter paper, putting the lid on the filter, and attaching it to the vacuum for filtration. A little amount of the liquid was poured at a time while ensuring the pellet was not poured. The filter paper was replaced. A clear 50 ml Falcon tube was used to pour the extract into the tube. The method was then set up. One of the leads was changed to point to the lysed cell extract. The sample value was pressed to make it green. The volume was altered to account for the amount present. For comparison with an established ladder through SDS-PAGE, a 2x loading dye and precision-plus protein unstained standards were utilized (Yan *et al,* 2016). Pellets were used for the ladder, liquid supernatant, and as an example sample: T2, T6, T12, T13, T14, T16, T17, and T18.

After this, a PCR tube and yellow micropipette were gotten, and 10 microliters of loading dye were placed into each. The sample was placed into each PCR tube as well (10 microliters). The tubes were labeled. The PCR tubes were placed into the Thermal Cycler and incubated (98 degrees C for 5 min). A non-native gel was used. The covering was placed to the gasket side and the colors were matched to the master position. An SDS buffer was poured into the gel to fill it up. A p1000 micropipette was taken and special gel micropipette tips were used to rest the buffer between the two. 5 microliters of ladder were needed, and 10 microliters of sample was put into each column in the gel. Samples were placed asymmetrically through the gel. When the wells were filled up, they were run at a current value of 150V. When the SDS-PAGE was read, the covers were removed and the gel was taken out of its covering. It was placed into the gel reader tray (where distilled water was poured to remove bubbles in the gel). The green button was pushed on the gel reader and opened on the gel reading computer software. The gel was labeled based on the positions of each respective sample. It was tossed in the disposal once the file had been saved on the SDS-PAGE gel reader system.

### ATPase assays with drug testing

**Figure 1:**
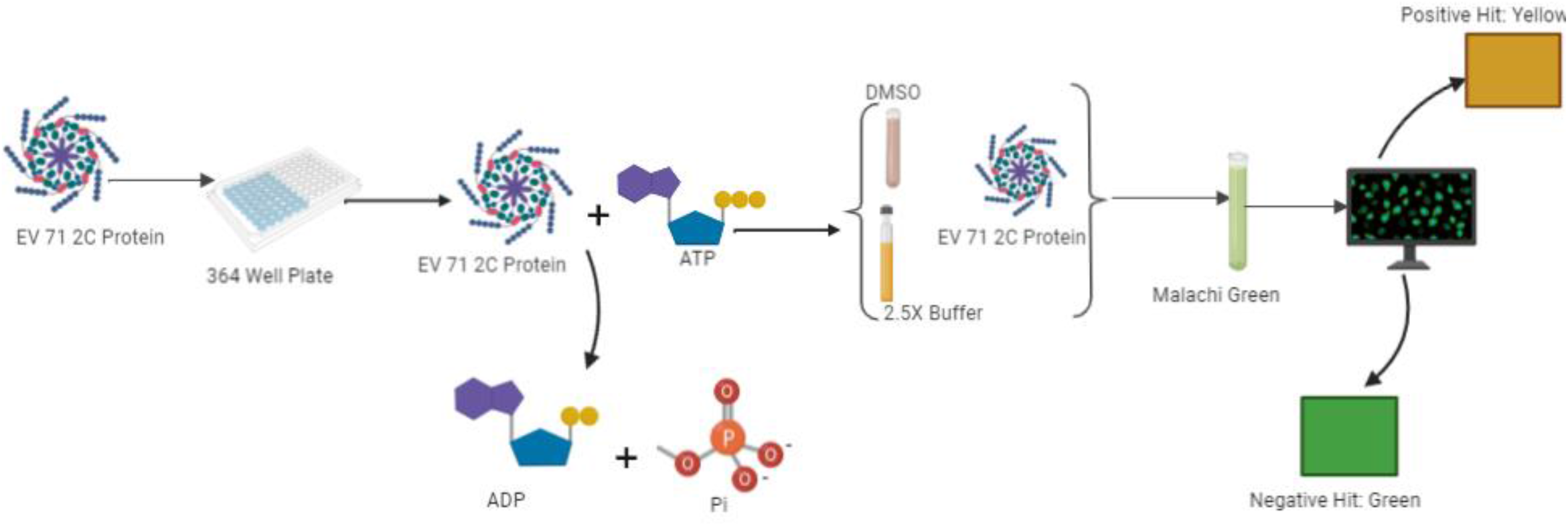
Mechanism of ATPase Assay.

**Figure 2:**
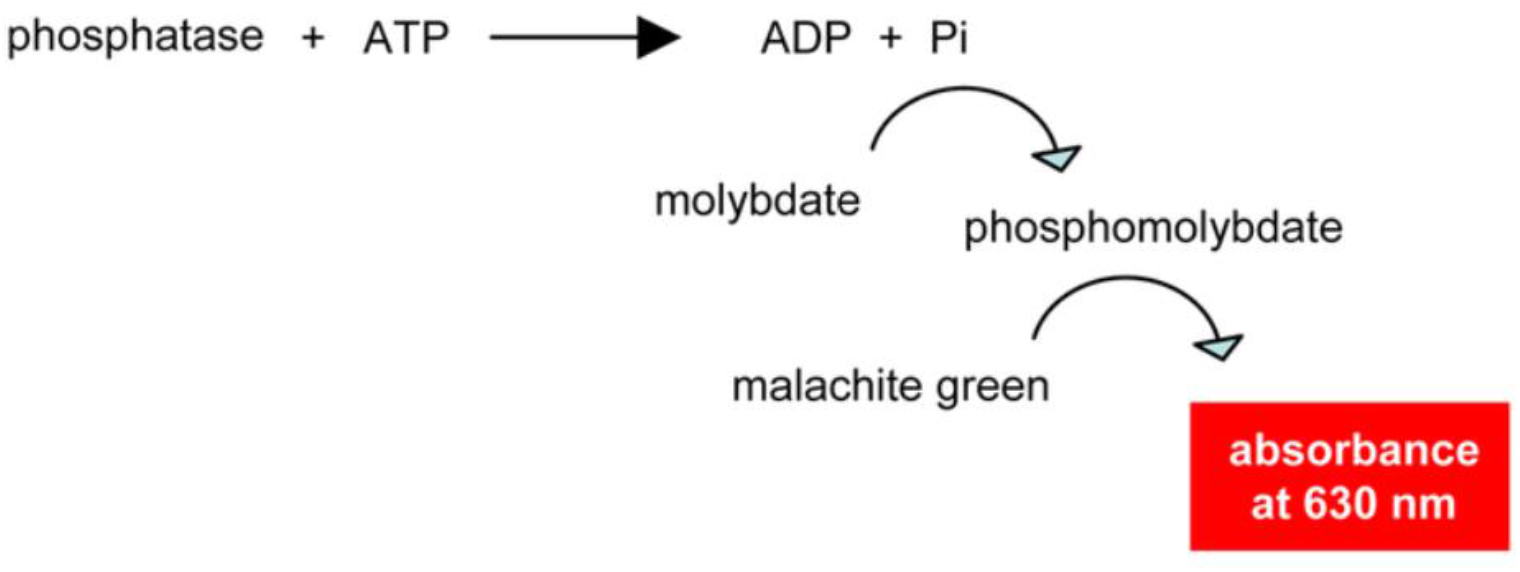
Depiction of the positive hit from ATPase assay. The plate reader would show a clear absorbance difference with a positive hit (Rompel *et al,* 2018).

The ATPase assay was done for the purpose of detecting the ATPase activity of the 2C protein. By detecting which compounds were able to stop the activity of the protein, it was useful for the development of antivirals (Ni *et al,* 2012). For this, a 100ml bottle was wrapped in foil. 45mg of Malachite green was dissolved in 100ml of Milli-Q H2O. It was mixed overnight using a blue stirrer. It was then stored at 4 degrees C for a maximum time period of 1 week. It was placed in 180 ml of 4.2% ammonium molybdate in 4N HCL. The solution was then dissolved with 7.56 g of ammonium molybdate in 120 ml of Milli-Q H2O and added to 60ml of 12N HCL. The 2C protein sample in Buffer B was diluted in Buffer B of TALON. The dilution was calculated to make the fraction reach a final concentration of 20 micromolar. In a 50ml Falcon tube, 12.5ml ammonium molybdate stock was mixed with 37.5ml of Malachite Green. This was mixed for a time period of 25 minutes. In a new 50 ml Falcon tube, 1ml of 1% Tween stock was added. Then, the assay was set up in a clear 384 well plate. It was designated in the following method: The final volume inside each well was capped at 50 microliters before the color reagent Malachi Green was added, and this consisted of 25 microliters of 2C protein that has been serially diluted with 25 microliters of Buffer A from AKTA. It was ensured that the micropipette did not touch the interior of the wells to prevent any possible cross contamination (Sun, 2006). Next, 25 microliters of ATPase were added to each of the wells. Once this was done, 25 microliters of the color reagent Malachi Green were incorporated into each of the wells. The 384 well plate was incubated in the shaker for 2 hours at 37 degrees Celsius. The ATPase assay was then brought to a plate reader for image processing and fluorescence readings.

## Results

**Figure 3:**
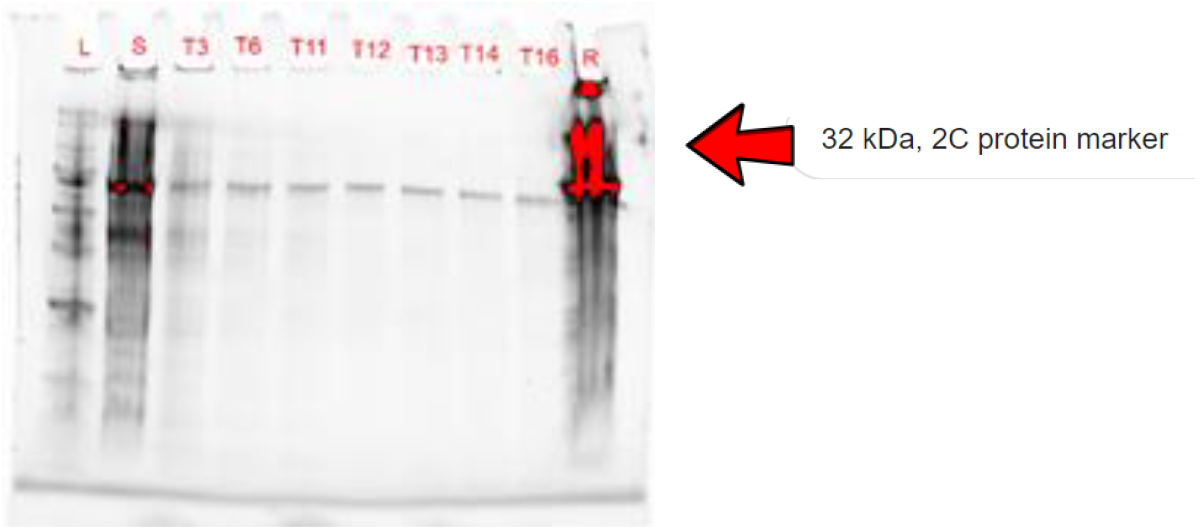
SDS-PAGE Gel after initial run via Nickel column on AKTA.

**Figure 4:**
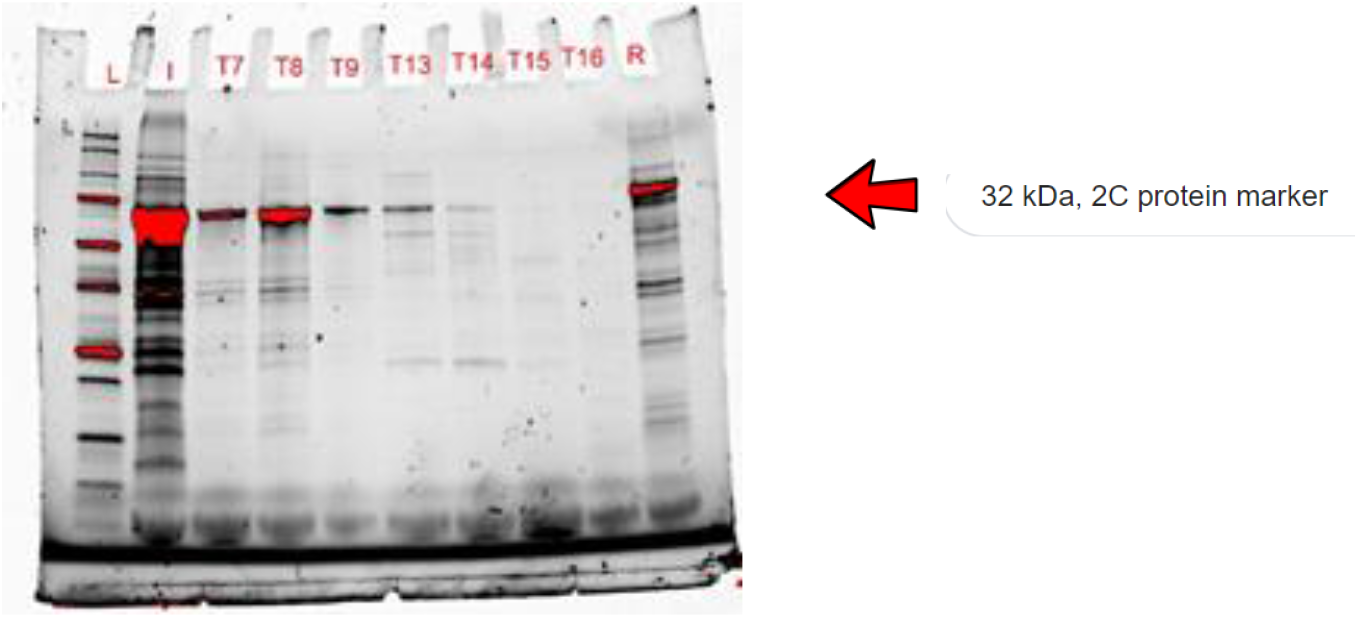
SDS-PAGE after SEC (Size Exclusion Chromatography). This protein concentration was used for the ATPase Assay.

**Figure 5:**
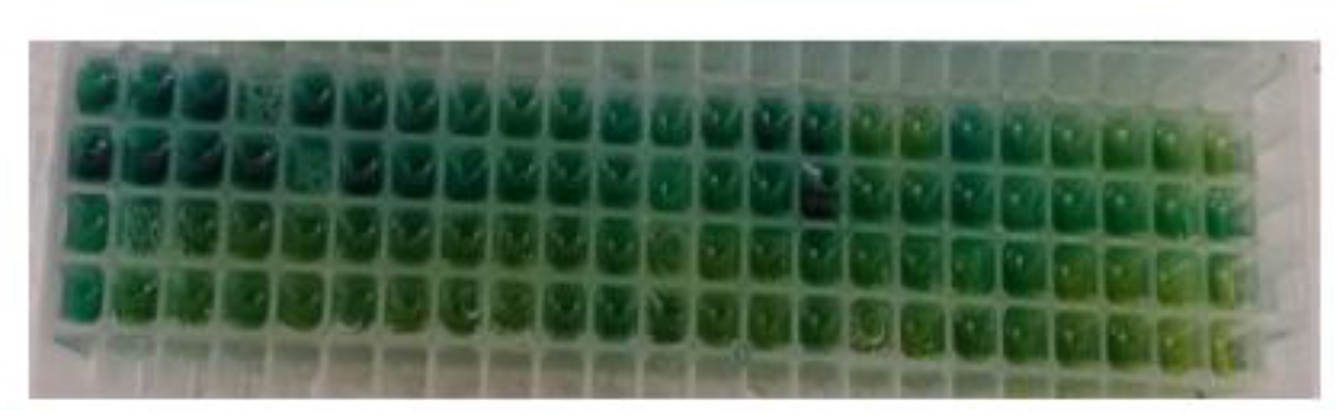
Setup of the ATPase assay.

**Table 1:** ATPase Assay average fluorescence values for each row.

| Dilution Factor of 2C protein present in each row (in micromolar) |  |  |  |  |  | Measure of Average Fluorescence in each row |
| --- | --- | --- | --- | --- | --- | --- |
| 1 |  |  |  |  |  | 0.9485 |
| 0.1 |  |  |  |  |  | 0.8437 |
| 0.01 |  |  |  |  |  | 0.9516 |
| 0.001 |  |  |  |  |  | 0.9419 |
| 0.0001 |  |  |  |  |  | 0.9154 |
| 0(control) |  |  |  |  |  | 0.5627 |

**Figure 6:**
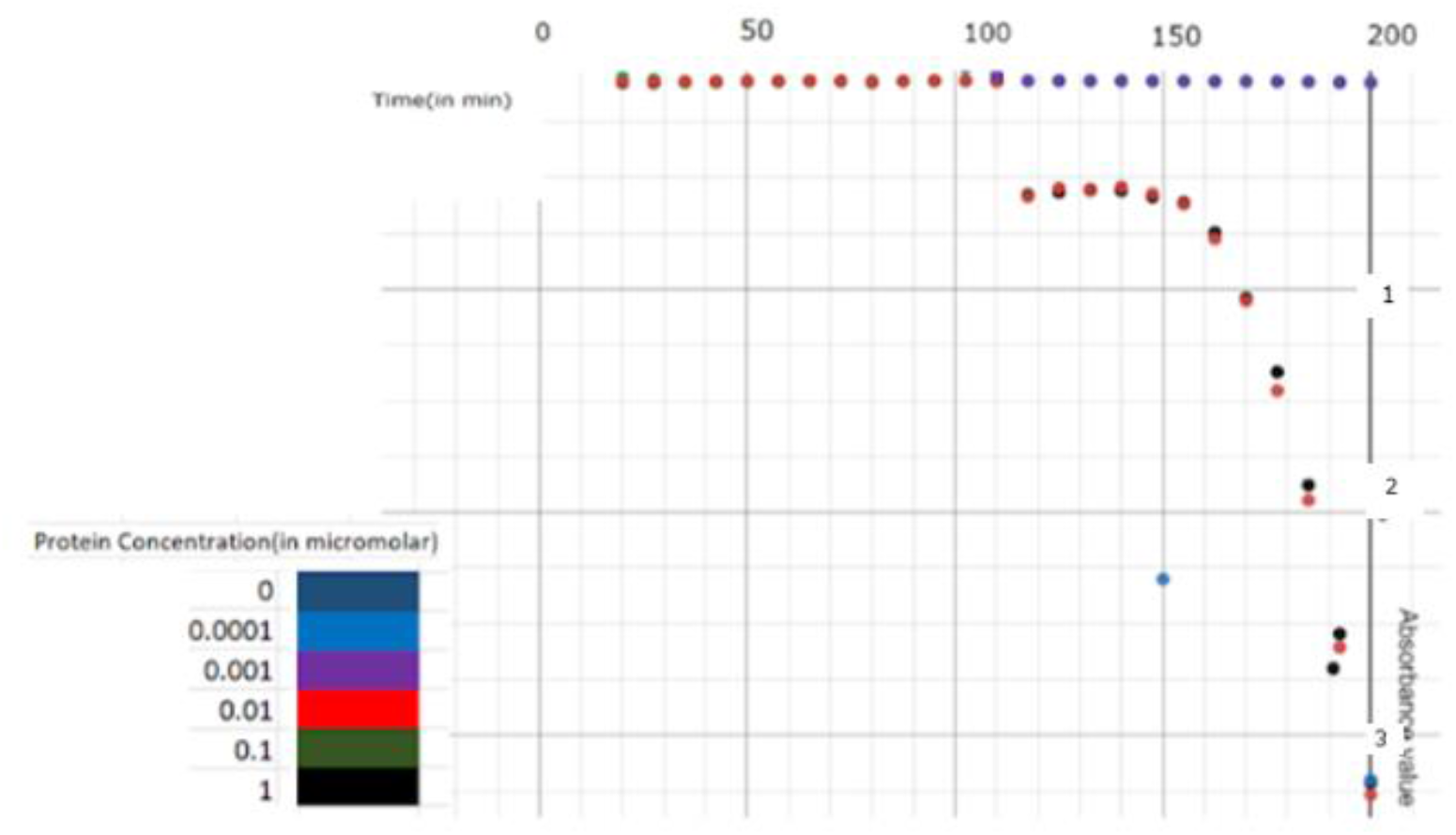
Scatterplot of individual ATPase fluorescence readings.

## Discussion

### Transformation of *E. coli* with the genetic information from Enterovirus 71

The results from the transformation showed that EV 71 genetic information was successfully transferred to the *E. coli* colonies. This result was obtained by physically transforming the *E. coli* through the method described above.

### Growth of the *E. coli* colonies in the lab setting

The results from the growth of the *E. coli* showed that they initially grew at an exponential rate without being hindered by resources or space. However, as time progressed, the rate of growth became logarithmic and slowly ceased. The growth of the bacteria colonies in the lab setting followed a normal growth rate pattern among *E. coli.*

### 2C protein purification

The results from the 2C protein purification showed that the protocol for extracting the protein was successful. The AKTA computer system was utilized to detect changes in conductivity at various time intervals as the sample was analyzed. These changes in conductivity were subjected to SDS-PAGE and compared to a standardized 2C protein ladder. The extracted protein and the ladder matched in terms of kDa length and band formation.

### ATPase assays with drug testing

The ATPase assays with drug testing revealed that a combination of two different antiviral compounds (Mitrofudil and N6-Benzyladenosine) was able to cease ATPase activity. This was qualitatively measured by the fluorescence exhibited by each well based on the presence of the color reagent. A yellow fluorescence indicates that there is no free phosphate available in the well, meaning that ATP was not being dephosphorylated. Four subsequent wells next to one another had to display this phenomenon to ensure that it was indeed the drug halting the breakdown of ATP. Mitrofudil and N6-Benzyladenosine together were able to prevent ATPase activity. A green fluorescence indicates that there is free phosphate available as a direct result of ATPase breaking down ATP to ADP and P. This shows that the other antiviral compounds would not inhibit ATPase activity and still allow new viral particle formation for this particular strain.

### Principle Results

The principle results from this experiment are that a conjunction of two antiviral compounds are best at blocking ATPase rather than each compound on its own. Coloration of wells when both drugs were combined appeared to be of a greater shade of yellow than when each compound was tested individually. This suggests that the mechanism of blocking ATPase is two-fold in EV 71.

### Limitations

It is important to note that more trials are needed to show that these two antiviral compounds are best suited for EV 71 treatment. These drugs have also not yet been tested on actual patients with Hand, Foot, and Mouth disease.

## Acknowledgements

This research would not have been possible without the assistance of Dr. Simon White and the equipment in his laboratory. These discoveries were all made in the Biophysics Building at the University of Connecticut, Storrs campus. This work was also partially funded by a grant awarded to the White Lab for advancements in virology.

## Notes

### Competing Interest Statement

The authors have declared no competing interest.

## References

1. Wong SS, Yip CC, Lau SK, Yuen KY. Human enterovirus 71 and hand, foot and mouth disease. Epidemiol Infect. 2010; 138:1071–89. [PubMed]

2. Ho M. Enterovirus 71: the virus, its infections and outbreaks. J Microbiol Immunol Infect. 2000; 33:205–16. [PubMed]

3. Centers for Disease Control and Prevention. Progress toward eradication of polio – worldwide, January 2011–March 2013. MMWR Morb Mortal Wkly Rep. 2013; 62:335–38. [PMC free article] [PubMed]

4. McMinn P, Lindsay K, Perera D, Chan HM, Chan KP, Cardosa MJ. Phylogenetic analysis of enterovirus 71 strains isolated during linked epidemics in Malaysia, Singapore, and Western Australia. J Virol. 2001; 75:7732–8. [PMC free article] [PubMed]

5. Li L, He Y, Yang H, Zhu J, Xu X, Dong J, et al. Genetic characteristics of human enterovirus 71 and coxsackievirus A16 circulating from 1999 to 2004 in Shenzhen, People’s Republic of China. J Clin Microbiol. 2005; 43:3835–9. [PMC free article] [PubMed]

6. Herrero LJ, Lee CS, Hurrelbrink RJ, Chua BH, Chua KB, McMinn PC. Molecular epidemiology of enterovirus 71 in peninsular Malaysia, 1997–2000. Arch Virol. 2003; 148:1369–5. [PubMed]

7. Chan KP, Goh KT, Chong CY, Teo ES, Lau G, Ling AE. Epidemic hand, foot and mouth disease caused by human enterovirus 71, Singapore. Emerg Infect Dis. 2003; 9:78–85. [PMC free article] [PubMed]

8. Wang JR, Tuan YC, Tsai HP, Yan JJ, Liu CC, Su IJ. Change of major genotype of enterovirus 71 in outbreaks of hand–foot-and-mouth disease in Taiwan between 1998 and 2000. J Clin Microbiol. 2002; 40:10–5. [PMC free article] [PubMed]

9. Chatproedprai S, Theanboonlers A, Korkong S, Thongmee C, Wananukul S, Poovorawan Y. Clinical and molecular characterization of hand–foot-and-mouth disease in Thailand, 2008– 2009. Jpn J Infect Dis. 2010; 63:229–33. [PubMed]

10. Tu PV, Thao NT, Perera D, Huu TK, Tien NT, Thuong TC, et al. Epidemiologic and virologic investigation of hand, foot, and mouth disease, southern Vietnam, 2005. Emerg Infect Dis. 2007; 13:1733–41. [PMC free article] [PubMed]

11. Zhang Y, Zhu Z, Yang WZ, Ren J, Tan XJ, Wang Y, et al. An emerging recombinant human enterovirus 71 responsible for the 2008 outbreak of hand foot and mouth disease in Fuyang city of China. Virol J. 2010;7:94. [PMC free article] [PubMed]

12. Sun LM, Zheng HY, Zheng HZ, Guo X, He JF, Guan DW, et al. An enterovirus 71 epidemic in Guangdong Province of China, 2008: epidemiological, clinical, and virogenic manifestations. Jpn J Infect Dis. 2011; 64:13–8. [PubMed]

13. Yang F, Zhang T, Hu Y, Wang X, Du J, Li Y, et al. Survey of enterovirus infections from hand, foot and mouth disease outbreak in China, 2009. Virol J. 2011; 8:508. [PMC free article] [PubMed]

14. Yan XF, Gao S, Xia JF, Ye R, Yu H, Long JE. Epidemic characteristics of hand, foot, and mouth disease in Shanghai from 2009 to 2010: enterovirus 71 subgenotype C4 as the primary causative agent and a high incidence of mixed infections with coxsackievirus A16. Scand J Infect Dis. 2012; 44:297–305. [PubMed]

15. Liu MY, Liu W, Luo J, Liu Y, Zhu Y, Berman H, et al. Characterization of an outbreak of hand, foot, and mouth disease in Nanchang, China in 2010. PLoS One. 2011;6: e25287. [PMC free article] [PubMed]

16. Ni H, Yi B, Yin J, Fang T, He T, Du Y, et al. Epidemiology and etiological characteristics of hand, foot, and mouth disease in Ningbo, China, 2008–2011. J Clin Virol. 2012; 54:342–8. [PubMed]

17. Tan X, Huang X, Zhu S, Chen H, Yu Q, Wang H, et al. The persistent circulation of enterovirus 71 in People’s Republic of China: causing emerging nationwide epidemics since 2008. PLoS One. 2011;6: e25662. [PMC free article] [PubMed]

